# CharPlant: A *De Novo* Open Chromatin Region (OCR) Prediction Tool for Plant Genomes

**DOI:** 10.1101/2020.10.27.358218

**Authors:** Yin Shen, Ling-Ling Chen, Junxiang Gao

**Affiliations:** Hubei Key Laboratory of Agricultural Bioinformatics, College of Informatics, Huazhong Agricultural University, Wuhan 430070, P. R. China; National Key Laboratory of Crop Genetic Improvement, Huazhong Agricultural University, Wuhan 430070, P. R. China

**Keywords:** Open chromatin region, Chromatin accessibility, Convolutional neural network, *De novo* prediction, Plant genome

## Abstract

Chromatin accessibility is a highly informative structural feature for understanding gene transcription regulation because it indicates the degree to which nuclear macromolecules such as proteins and RNA can access chromosomal DNA. Studies show that chromatin accessibility is highly dynamic during stress response, stimulus response, and developmental transition. Moreover, physical access to chromosomal DNA in eukaryotes is highly cell-specific. Therefore, current technologies such as DNase-seq, ATAC-seq, and FAIRE-seq reveal only a portion of the open chromatin regions (OCRs) present in a given species. Thus, the genome-wide distribution of OCRs remains unknown. In this study, we developed a bioinformatics tool called CharPlant for the *de novo* prediction of chromatin accessible regions in plant genomes. To develop this tool, we constructed a three-layer convolutional neural network (CNN) and subsequently trained the CNN using DNase-seq and ATAC-seq datasets of four plant species. The model simultaneously learns the sequence motifs and regulatory logics, which are jointly used to determine DNA accessibility. All of these steps are integrated into CharPlant, which can be run using a simple command line. The results of data analysis using CharPlant in this study demonstrate its prediction power and computational efficiency. To our knowledge, CharPlant is the first *de novo* prediction tool that can identify potential OCRs in the whole genome. The source code of CharPlant and supporting files are freely downloadable from https://github.com/Yin-Shen/CharPlant.

## 1 Introduction

In eukaryotic genomes, most of the chromatin is tightly coiled in the nucleus, but some regions, known as open chromatin regions (OCRs) or chromatin accessible regions, are loosely formed after chromatin remodeling. Whether the chromatin is loosely or tightly coiled largely determines transcriptional regulation [1,2]. A number of cis-regulatory elements interact with trans-acting factors for transcriptional regulation, and cis-trans elements with regulatory functions participate in the process of transcriptional regulation by binding to OCRs [3,4]. For example, when a transcription factor binds to an OCR, it recruits other proteins to initiate the transcription of nearby genes. Therefore, a complete genome-wide map of potential open chromatin loci is helpful for the investigation of changes in the nucleosome location and for the discovery of genome regulatory elements and gene regulatory mechanisms [5,6]. Chromatin accessibility information has even been proven to be valuable for the early diagnosis and treatment of cancer [7,8].

The open regions of chromatin are easier to excise than other regions. Therefore, researchers often use enzymes, such as nuclease and transposase, or physical methods to digest the chromatin. The cleavage-sensitive sites are then sequenced using various technologies, such as DNase I hypersensitive sites sequencing (DNase-seq), assay for transposase accessible chromatin sequencing (ATAC-seq), and formaldehyde-assisted isolation of regulatory elements sequencing (FAIRE-seq), to obtain further information. DNase-seq has been used for a long time; however, it requires a large amount of starting material (~10^7^ cells). On the other hand, ATAC-seq requires a significantly smaller sample (<10^5^ cells) and has the advantage of requiring no antibody. Therefore, ATAC-seq has become the method of choice in recent years [9,10]. However, none of these techniques have been able to solve the problem of open chromatin determination. All nuclease-based methods exhibit a preference for specific sequences for cleavage, depending on the nuclease, which is a major flaw. For example, DNase I exhibits a strong preference for specific sequences, and many DNase-seq results reflect cleavage preferences rather than actual protein binding [3,11,12]. Similarly, in the ATAC-seq method, a preference for cleavage sites has been observed with some of the Tn5 enzymes, resulting in “false DNA footprints” [13]. Although these technologies have been commonly used in human and animal studies [14], their application in plants is still in the exploratory stage. This is because of structural differences between plant and animal cells. Unlike animal cells, plant cells possess a cell wall and numerous chloroplasts, mitochondria, and other organelles that contaminate the assay. Consequently, OCR data have been obtained using DNase-seq and ATAC-seq only in a small number of model plant species, including *Oryza sativa* [15], *Arabidopsis thaliana*, *Medicago truncatula*, *Solanum lycopersicum* [16], and *Hordeum vulgare* [17].

Previous studies show that chromatin accessibility is highly dynamic rather than static. OCRs usually change during stress response, stimulus response, and developmental transition [18,19]. Moreover, OCRs in different species are significantly cell-specific [4]. More than 40% of the OCRs in human T-cells differ between functional and exhausted cells at different time points [20]. Chromatin accessibility also varies considerably among different cells in *Drosophila melanogaster*, *A. thaliana*, and *O. sativa* [16,21]. Consequently, the current DNase-seq, ATAC-seq, and FAIRE-seq data represent only some of the OCRs and do not present the entire chromatin accessibility information about a given species [22,23]. Thus, a global overview of the distribution of OCRs in genomes is lacking. Moreover, these experimental technologies are generally expensive and time-consuming [3].

Proteins recognize specific motifs and epigenetic modifications of the DNA sequence that influence its accessibility [24]. After training on a specific dataset, machine-learning algorithms can collect sequence information and predict protein-binding sites, DNA accessibility, histone modifications, and DNA methylation patterns. Many algorithms have been developed to predict regulatory elements, such as Basset [25], Deeperdeepsea [26], DeepBind [27], and DeepCpG [28]. However, these algorithms have a few limitations. First, most of the algorithms are not designed for the prediction of OCRs; instead, they are designed for the prediction of 1) regulatory fragments that bind to transcription factors or RNA-binding proteins, 2) DNase sensitivity, and 3) genomic variants. Second, almost all previous studies using such algorithms are based on human or mouse data; however, the OCRs of plants and animals exhibit significantly different characteristics. For example, approximately 39% of the DNase I hypersensitive sites (DHSs) are associated with introns in the human genome, which is remarkably higher than the proportion of intron-associated DHSs in *O. sativa* (11%) and *A. thaliana* (5%) [29]. Third, most of the existing methods are developed as conventional classifiers that classify sequence fragments of a certain length (hundreds of base-pairs) as regulatory regions, instead of scanning the whole genome.

OCRs are usually rich in various elements and specific motif-binding factors. Therefore, it is feasible to scan the genome and predict chromatin accessibility by learning the motifs and their distribution from OCR data. Here, we developed a *de novo* OCR prediction tool called CharPlant (<u>C</u>hromatin <u>A</u>ccessible <u>R</u>egions for <u>P</u>lant), based on deep learning, to provide a genome-wide overview of OCRs in a given plant species. We constructed training datasets using the DNase-seq and ATAC-seq data of four plant species. CharPlant simultaneously learns the relevant sequence motifs and regulatory logics, which are jointly used to determine DNA accessibility. The trained model accepts DNA sequences or scaffolds as input and generates an outline of OCRs in a .bed file as output (Figure 1A). To our knowledge, this is the first tool capable of *de novo* prediction of OCRs from the DNA sequence.

**Figure 1.**
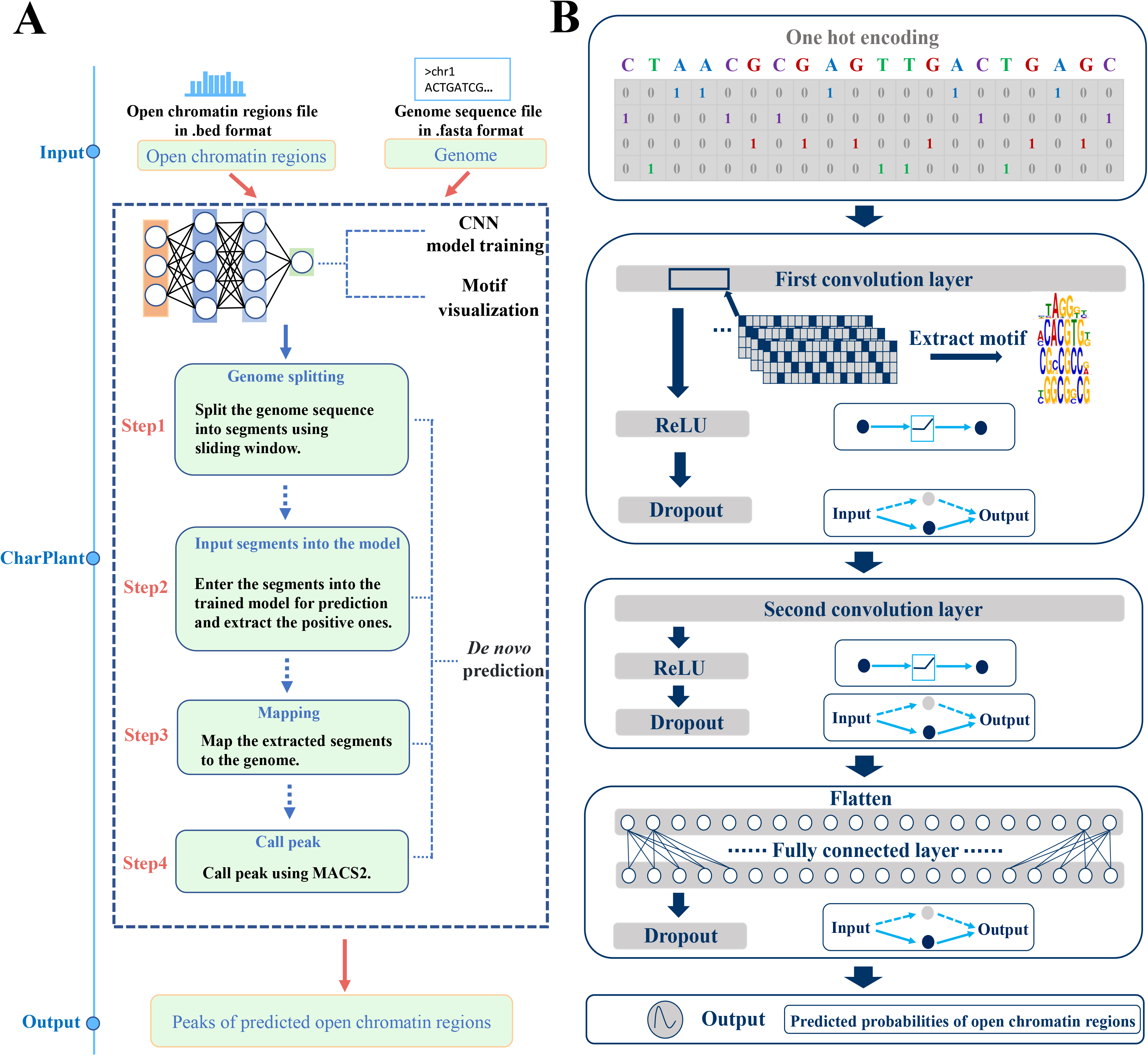
Steps involved in the construction and execution of CharPlant, a *de novo* OCR prediction tool. **A**. *De novo* OCR prediction pipeline. **B**. Construction and training of the CharPlant network. CNN, convolutional neural network; OCR, open chromatin region.

## 2 Materials and methods

### 2.1 Construction of datasets

Considering that a dataset has to be constructed using representative plant species that have both OCR assay data and high-quality reference genomes, the DNase-seq data of *O. sativa* and the ATAC-seq data of four plant species (*A. thaliana*, *S. lycopersicum*, *M. truncatula*, and *O. sativa*) were downloaded from the PlantDHS database at http://plantdhs.org and the GEO of NCBI (GEO Accessions: GSE101482 and GSE75794) at https://www.ncbi.nlm.nih.gov/geo [15], respectively. Detailed information about the DNase-seq and ATAC-seq data is listed in Table 1, and basic information about the reference genomes of these four plant species is listed in Table 2. Because the DNase-seq and ATAC-seq data used here represent diverse plant species of both dicot and monocot lineages and different cell types of the same species (*O. sativa*), this model applies to a broad range of plant species with distant evolutionary relationships. The use of two mainstream technologies, DNase-seq and ATAC-seq, allowed the inclusion of nuclease-based and transposase-based data. MACS2 software was used for peak calling with default parameters [30]. Because of higher statistical power using long fragments, peaks longer than 200 bp were filtered out as positive samples. Positive sequences were shuffled to generate the negative dataset with fasta-shuffle-letters in the MEME software [31]. Unlike the random interception of fragments from the DNA sequence for use as negative samples, shuffling ensures that the negative and positive samples have identical composition of all four bases [27]. Shuffling also maintains a balance between positive and negative samples when constructing the dataset. In an unbalanced dataset, classification algorithms focus on the class containing the most samples, which degrades the classification performance of the class that contains a small number of samples. Most machine-learning algorithms do not work well with unbalanced datasets. Therefore, to construct the dataset in this study, a negative sequence was generated using each positive sequence, i.e., the number of positive and negative samples were equal in number. The samples were divided into three sets, training set, validation set, and test set, which accounted for 60%, 20%, and 20% of the data, respectively. The sample numbers of the three sets are listed in Table 1.

**Table 1.** Detailed information about DNase-seq and ATAC-seq datasets.

| Species | Data type | Database | Accession number | Treatment | Sample number |  |  |
| --- | --- | --- | --- | --- | --- | --- | --- |
|  |  |  |  |  | Training set | Validation set | Test set |
| *O. sativa* | DNase-seq | PlantDHS | NA | Seedling and callus | 70,810 | 23,604 | 23,604 |
| *O. sativa* | ATAC-seq | NCBI | GSE101482 | Root of 7-day-old seedlings;<br>two biological replicates | All GSE101482 data<br>were used as the test set. |  | 39,868 |
| *O. sativa* | ATAC-seq | NCBI | GSE75794 | 2<sup>nd</sup> leaf of 14-day-old seedlings; heat stress<br>and recovery, dehydration stress and<br>recovery | All GSE75794 data<br>were used as the test set. |  | 30,634 |
| *A. thaliana* | ATAC-seq | NCBI | GSE101482 | Root cells; two biological replicates | 13,778 | 4,594 | 4,596 |
| *S. lycopersicum* | ATAC-seq | NCBI | GSE101482 | Root cells; two biological replicates | 32,168 | 10,724 | 10,726 |
| *M. truncatula* | ATAC-seq | NCBI | GSE101482 | Root cells; two biological replicates | 27,982 | 9,328 | 9,330 |

**Table 2.**
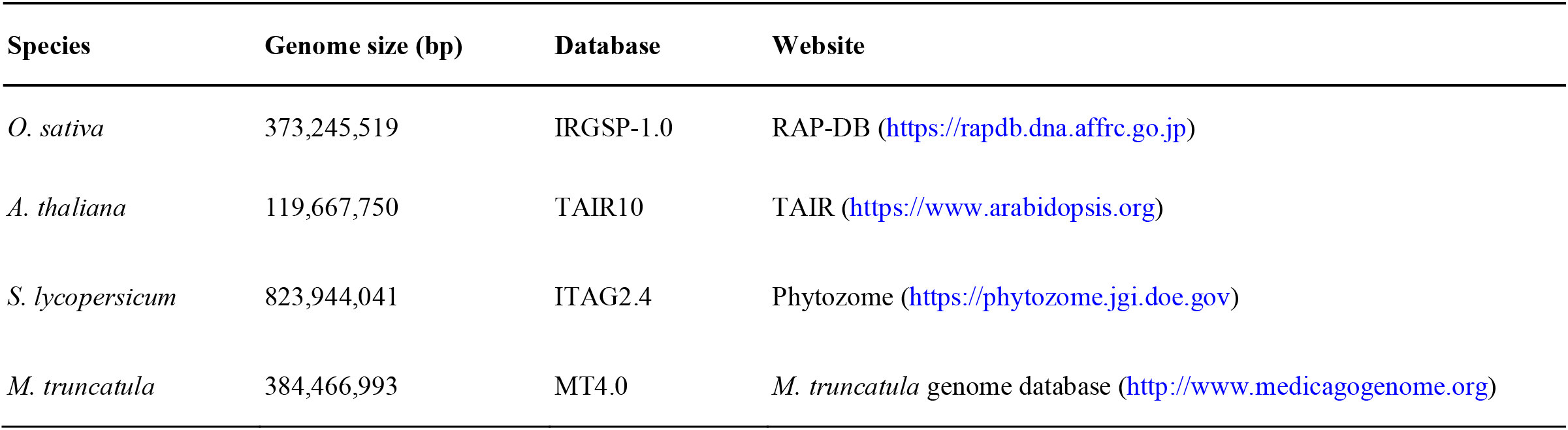
Genome-related information on the four plant species used in this study.

| Species | Genome size (bp) | Database | Website |
| --- | --- | --- | --- |
| *O. sativa* | 373,245,519 | IRGSP-1.0 | RAP-DB (https://rapdb.dna.affrc.go.jp) |
| *A. thaliana* | 119,667,750 | TAIR10 | TAIR (https://www.arabidopsis.org) |
| *S. lycopersicum* | 823,944,041 | ITAG2.4 | Phytozome (https://phytozome.jgi.doe.gov) |
| *M. truncatula* | 384,466,993 | MT4.0 | *M. truncatula* genome database (http://www.medicagogenome.org) |

### 2.2 Construction of the CharPlant model

The CharPlant model was trained using ATAC-seq or DNase-seq data of four plants, and each plant had its own model parameters. The model was based on a multilayer convolutional neural network (CNN; Figure 1B). The CNN model, which originated from artificial neural networks, contains perceptrons with multiple hidden layers and combines low-level features to form more abstract high-level attributes or features for discovering feature representations of data [32,33]. The motivation is to build a neural network that can simulate the human neuron for analysis and learning, and imitate the mechanism of the human brain to interpret data, such as images, sounds, and texts [34–36]. Unlike traditional methods, in which features are manually selected in the pre-processing stage, the CNN adaptively extracts features from large-scale training datasets. It then maps input data to high-dimensional representations with abundant information by nonlinear transformation, thus simplifying classification or regression. Early application of the CNN model in DNA sequence analysis surpasses existing mature algorithms, such as support vector machines or Random forests, in predicting protein binding and DNA sequence accessibility [25,27].

To achieve high computational efficiency, our model was designed with only three hidden layers; the first and second layers were convolutional, whereas the third layer was fully connected. The CNN model used in CharPlant requires binary vectors as input. Each input DNA fragment is first converted into a 4 × *n* matrix, where *n* represents the length of the input fragment. Thus, each base is preprocessed with “one-hot” encoding (A: [1, 0, 0, 0]; C: [0, 1, 0, 0]; G: [0, 0, 1, 0]; T: [0, 0, 0, 1]; N: [0, 0, 0, 0]), and the sequence is converted into a matrix with four columns. The first layer of the CNN model contains convolutional filters for the identification of low-level features in a given DNA sequence. A convolutional filter is essentially a motif prober that scans each input matrix to discover potential patterns. The identification of low-level DNA features involves the following steps. First, each input sequence is fed into the first convolutional layer, and the convolution kernel slides over the sequence fragment to calculate the activation score. If the activation score of the convolution kernel at a certain position is greater than the preset threshold, the sequence segment centered at that position will be identified and represented by the position frequency matrix (PFM) of four base frequencies. Then, the PFM is used to calculate the information entropy and is transformed into position weight matrix (PWM), which is widely used for the representation of motifs. The PMW contains four rows, and describes the entropy of four bases at each position [25,37]. Subsequently, the sequence logo is used to visualize the motif, i.e., the base size of each position indicates the possibility of the base at this position. To obtain the activation score of the convolutional filter, the rectified linear unit (ReLU) was used as the activation function for three hidden layers. The ReLU function f(x) is calculated as follows:

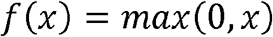

where *x* is the input.

The ReLU function is used by neurons just like traditional activation functions such as sigmoid or hyperbolic tangent. Compared with the conventional activation function, ReLU has much less computational complexity for calculating the error gradient in back propagation. Additionally, when the conventional activation function propagates backward, it is likely that the derivative will approach zero, which would make it impossible to complete the training of deep network. However, ReLU overcomes this shortcoming very well.

To overcome the problem of over-fitting, random dropout was set after every hidden layer in the model. Dropout is an optimization method for resolving over-fitting and gradient disappearance in deep neural networks. In the learning process of the neural network, the weight of randomly selected nodes in the hidden layer was set at zero. Because different nodes were reset to zero after each iteration, the importance of each node was balanced. Because of the use of random dropout, each node of the neural network contributed roughly equally to the training, and there was no case where a few high-weight nodes completely dominated the output. In this study, the dropout probability was set at 0.6, i.e., the weights of 60% of the neurons were set to zero in every iteration.

The architecture of the second layer was the same as that of the first layer and was based on three key technologies: convolutional network, ReLU activation function, and dropout. The second layer combined low-level motif features to form abstract high-level attributes. In the CNN model, a fully connected layer was set behind the two convolutional layers. Each neuron in the fully connected layer was connected with all the neurons in its preceding layer to integrate local sequence information with class discrimination in the convolutional layer. The fully connected layer contained 200 neurons, and it flattened the matrix into a column vector. The weights of links were calculated by a linear regression algorithm, but linear regression could only predict the continuous value, which did not solve the classification problem. Therefore, the output layer determined whether the input sequence belonged to the positive or negative class, depending on the calculations. The final output layer used sigmoid function to perform nonlinear transformation and mapped the results of the fully connected layer from (−∞, +∞) to (0, 1), which indicated the probability of open chromatin sequence. The goal of the training model was to minimize the error between predicted and labeled values, i.e., to minimize the cost function. A series of cost functions is available. Here, the binary cross-entropy cost function was used because it can overcome the problem of gradient disappearance when calculating gradient descent, thus showing high learning efficiency.

### 2.3 Implementation of CharPlant

CharPlant, implemented in Python, was based on TensorFlow 1.2.0, an open source machine-learning platform developed by Google. Additionally, a widely used workflow management system, Snakemake, was employed to combine a series of steps into a single pipeline that can be run by an inexperienced user using simple command line entries [38]. Steps in the workflow are described in terms of the rules defined using the input and output and Shell and Python codes. The workflow determines the steps that need to be performed and produces one or more output files. Dependencies between rules were automatically resolved, and rules were automatically parallelized when possible. A text file titled “Snakefile” was created, which defined the input and how the output was created from the input. CharPlant can learn OCR features from DNase-seq or ATAC-seq data and predict potential chromatin accessible regions in a plant genome *de novo*. It performs four steps: (i) data pre-processing, (ii) model training, (iii) motif visualization, and (iv) *de novo* prediction. If all the steps are successful, CharPlant outputs the results of the predicted OCRs in a .bed file in the directory CharPlant/peak.

Snakefile has a number of parameters such as epoch number, learning rate, batch size, and dropout, which could be adjusted using a configuration file (config.yaml). The “config.yaml” file used in this study provides default values and their meaning for each parameter. It is not necessary for the users to modify the default values, except for two directories as shown below. For example, the parameter “genome” represents the input genome file in .fasta format, and the parameter “bed” represents the output OCRs file in .bed format. The user would need to replace “Yourpath” with the true path in which CharPlant was installed.

genome: Yourpath/CharPlant/example/oryza_sativa.fa

bed: Yourpath/CharPlant/example/oryza_sativa.bed

### 2.4 Installation and execution of CharPlant

CharPlant is currently available for Linux-based operating systems. To install and run CharPlant, download the package from the GitHub development platform at https://github.com/Yin-Shen/CharPlant and then set “CharPlant” as the current directory. The subdirectory CharPlant/example contains the reference genome (file oryza_sativa.fa) and DNase-seq data of *O. sativa* as an example (ory_whole.bed). All Python and Shell scripts are in the subdirectory CharPlant/src. Some fundamental Python packages, such as numpy, matplotlib, and keras, will be needed for scientific computing and network construction. Supplementary material provides a detailed CharPlant manual, installation steps, and parameter settings for the abovementioned Python packages, and a complete “config.yaml” and “Snakemake” file. Users can run the program by typing the following command:

$ CharPlant.sh

## 3 Results and Discussion

### 3.1 Motifs identified by CharPlant

The positive dataset was obtained from the peaks of ATAC-seq and DNase-seq data, and the negative samples were generated by shuffling the positive samples, as described above. In the positive samples, motifs are usually clustered for protein binding, whereas the negative samples generally have far fewer motifs. A DNA motif is defined as a short similar recurring pattern of nucleotides, with many biological functions. A previous study showed that sequence motifs are roughly constant in length, and are often repeated and conserved [39]. Based on the difference between positive and negative datasets, our model learned the sequence motifs and the regulatory logics with which they are combined to determine DNA accessibility. The convolutional layer searched for the motifs along the genome sequence and produced a matrix, with rows representing neurons and columns representing positions. Determination of OCRs was based on the accurate identification of motifs. We compared the motifs learned by the convolution kernels with the known motifs in the JASPAR database [37]. The results showed that many of the motifs predicted by our model were previously known and experimentally validated. For example, JASPAR has eight known motifs in *O. sativa*, of which six were identified by our model (Figure 2A). We also identified many known motifs of other plants, some of which are shown in Figure 2B. Notably, sequence motifs were very small in size (6–19 bp), whereas intergenic regions were very long and highly variable, thus making motif discovery a very difficult task. Therefore, the number of motifs in plant genomes and their positions with respect to target genes remain unclear. Given that JASPAR has a very limited repertoire of only 501 experimentally validated motifs, some of the identified sequences not included in the database could be potential motifs, which might be experimentally validated in the future. Overall, CharPlant can detect motifs of various lengths, which can be subsequently used for the identification of OCRs.

**Figure 2.**
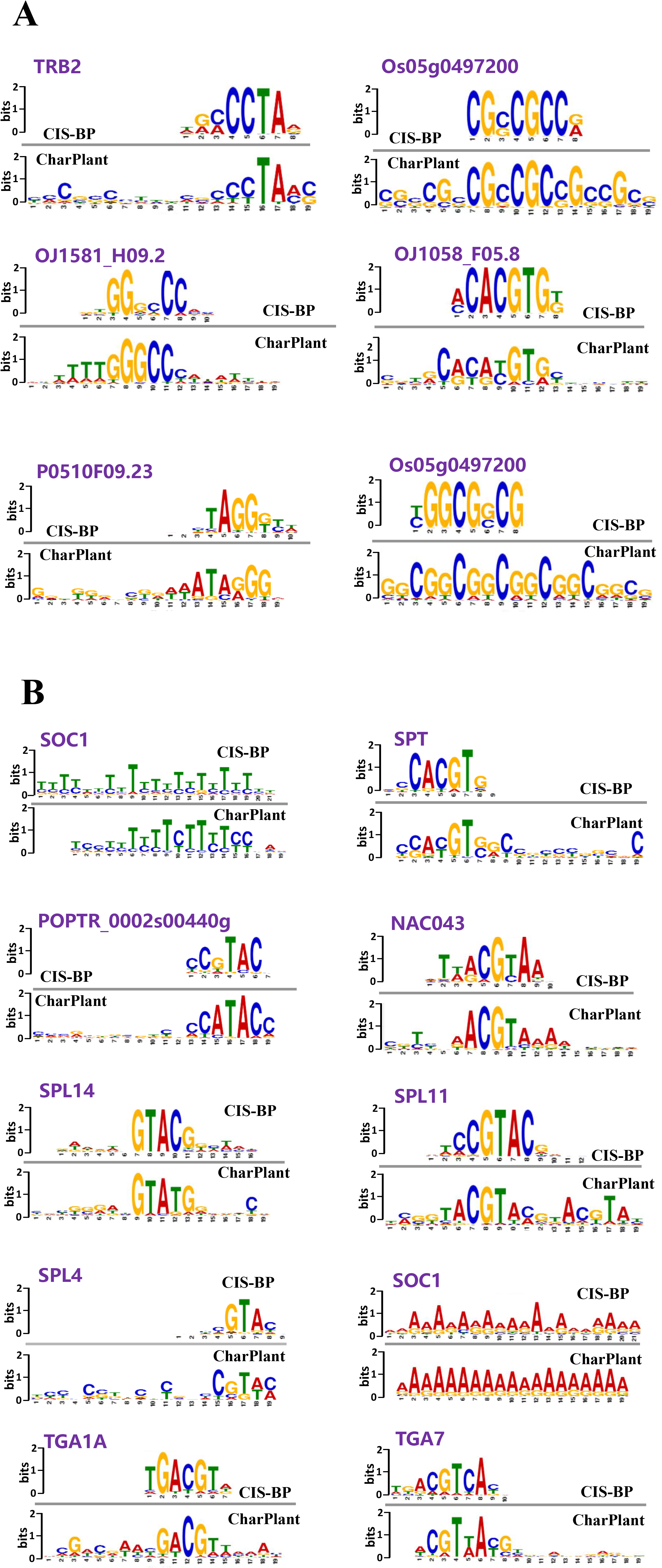
Motifs identified by CharPlant in *Oryza sativa* and other plant species. **A**. Six of the eight known *Oryza sativa* motifs in the JASPAR database identified by CharPlant. **B**. Motifs of other plant species in JASPAR identified by CharPlant.

### 3.2 Performance comparison between CharPlant and other methods

CharPlant is designed as a *de novo* OCR prediction tool. By contrast, almost all current methods have been developed as conventional classifier algorithms and cannot scan the genome sequence to discover OCRs. Moreover, the architecture and parameters of these models have been developed based on human and animal data. Therefore, a strict comparison of these methods with CharPlant is difficult. Because it was impossible to compare *de novo* prediction with other methods, we compared the learning ability and computational efficiency of CharPlant with two state-of-the-art deep learning algorithms, Basset [25] and Deeperdeepsea [26], using chromatin accessibility plant data. Basset is an open source package and learns the functional activity of DNA sequences from genomic data. The authors applied Basset to a compendium of accessible genomic sites mapped in 164 cell types by DNase-seq and showed greater predictive accuracy than previous methods. We revised Basset to adapt it to plant data. Deeperdeepsea is a recently published PyTorch-based deep learning library for any biological sequence data. We downloaded the package from https://selene.flatironinstitute.org/. Each method was adjusted to its best state and trained using the dataset constructed in this study, as described above. We calculated the false-positive rate vs. true-positive rate to plot receiver operating curves (ROCs) and determined the area under the ROC curves (AUCs; Figure 3, Figure S1). In Figure 3A, the curves were obtained using the DNase-seq data of *O. sativa*. Although the AUCs of CharPlant were slightly better than those of Basset and Deeperdeepsea on the *O. sativa* DNase-seq data, no major difference was observed among the three methods. However, the performance of these three methods on the other three datasets was quite different. Although the ROC and AUC of Basset on *O. sativa* and *A. thaliana* datasets were similar to those of CharPlant, the prediction accuracy of Basset was only ~50% with *S. lycopersicum* and *M. truncatula* datasets. Thus, the results of Basset were equivalent to a random guess, indicating that Basset failed to predict OCRs. Similarly, Deeperdeepsea failed on the datasets of all plant species, except *O. sativa.* By contrast, our method could be applied to all datasets and achieve consistent performance. Basset and Deeperdeepsea are both excellent methods for the prediction of regulatory elements and have been proven to produce accurate results after training on human data. However, these methods do not work on plant datasets, as shown in this study. This is likely because the structural design and hyperparameter choice of the model are not suitable for plant datasets. The characteristics of DNA sequences differ greatly between plants and animals. To shift an algorithm from the animal to the plant system, replacing the animal training set with a plant dataset is not sufficient; instead, to achieve similar performance, it is often necessary to make substantial changes to the model structure, essentially transforming it into a new model.

**Figure 3.**
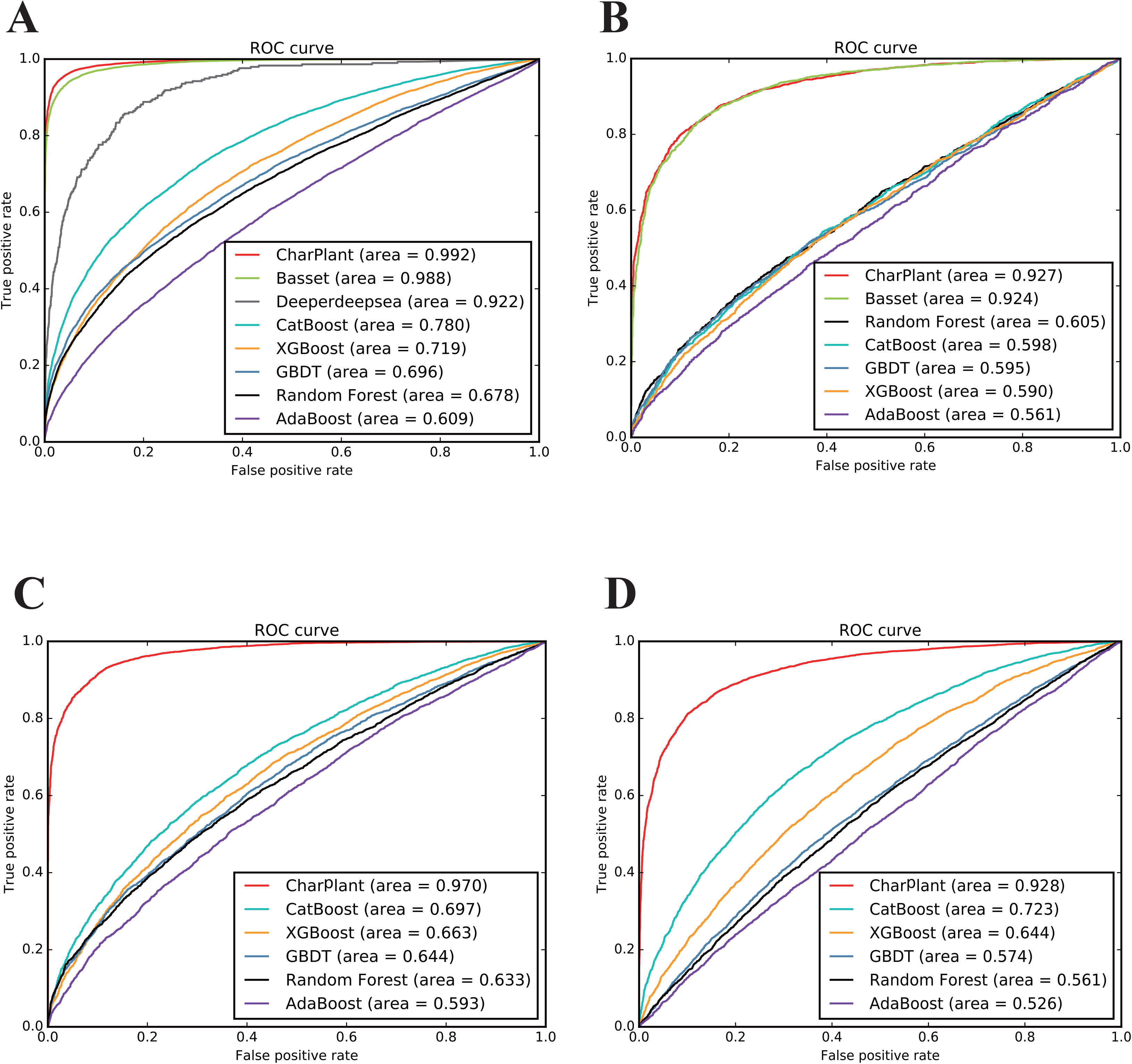
Comparison of ROCs and AUCs of CharPlant with those of comparative methods on four datasets. **A**. *Oryza sativa*. **B**. *Arabidopsis thaliana*. **C**. *Medicago truncatula*. **D**. *Solanum lycopersicum*. ROC, receiver operating curve; AUC, area under the ROC curve.

To further validate the performance of CharPlant, we compared our model with machine-learning methods including Random forest, Adaboost, GBDT, XGBboost, and CatBoost on all four plant datasets. These machine-learning methods were implemented using the Scikit-learn package, a widely used library that supports supervised and unsupervised learning [40]. We computed the precision and recall ratios of the comparative methods, and plotted the precision recall curves in Figure S2. Analysis of the precision recall curves of CharPlant, Basset, and Deeperdeepsea led us to a similar conclusion to that obtained from the analysis of ROCs described above (Figure 3). When the other comparative methods, including Random forest, Adaboost, GBDT, XGBboost, and CatBoost were used with the four plant datasets, their performance was similar, and their curves were close to each other. However, the performance of each of these algorithms was significantly inferior to that of neural network methods.

In some instances, the method of sample partitioning influences model evaluation. To avoid the randomness of a single training set and test set, we performed 10-fold cross validation. The dataset was divided into 10 parts and each part was used in turn as a test dataset, and nine were used as the training dataset. ROC and precision recall curve was plotted for each test (Figure S3). All samples were used as training and test sets, and each sample was tested one time. The majority of precision recall curves overlapped, indicating that precision and recall were stable when using different dataset partition methods. Similar conclusions could be drawn from ROCs.

To compare the computation efficiencies, we trained and tested CharPlant, Basset, and Deeperdeepsea on the central processing unit (CPU) and graphics processing unit (GPU). The manufacturers and models are as follows: Tesla P100-PCIE-16GB (GPU) and Intel(R) Xeon(R) Gold 6140 CPU @ 2.30 GHz (CPU). The comparison was performed on the DNase-seq dataset of *O. sativa*. The results showed that CharPlant took significantly less time than Basset and Deeperdeepsea on both CPU and GPU (Table 3).

**Table 3.** Training and testing times of the three methods on CPU and GPU.

| Method | Training time (s) |  | Testing time (s) |  |
| --- | --- | --- | --- | --- |
|  | GPU | CPU | GPU | CPU |
| **CharPlant** | 2,558 | 280,640 | 21 | 2,371 |
| **Basset** | 3,205 | 382,093 | 28 | 3,533 |
| **Deeperdeepsea** | 5,520 | 13,502,400 | 290 | 12,465 |

*Note:* cpu, central processing unit; gpu, graphics processing unit.

### 3.3 *De novo* prediction of OCRs in genomes

To enable the prediction of OCRs from long DNA sequences or complete genomes using CharPlant, we used the sliding-window method to split the sequence into fragments. The window width was set at 36 bp. Generally, a smaller sliding step is helpful for the accurate prediction of the location of OCRs; however, the computational complexity with a smaller sliding step is significantly higher than that with a large sliding window. To compromise the calculation efficiency and accuracy, the sliding step was set at 5 bp. The trained model was used to calculate the probability of chromatin accessibility of these fragments, and then the peaks of OCRs in these fragments were called using the MACS2 tool, with default parameters [30]. We scanned the whole genome sequences of four plant species using the CharPlant model and aligned the predicted OCRs with the DNase-seq or ATAC-seq dataset to validate the performance of CharPlant. The training datasets were obtained from a single cell type at a specific time, yet we tried to predict all potential OCRs in different tissues at different times. The results showed that the number of OCRs in the latter was higher than that in the former. Notably, the number of OCRs predicted by CharPlant in all datasets was much larger than that detected by DNase-seq or ATAC-seq assays (Figure 4). CharPlant predicted 153,594 potential OCRs in the *O. sativa* seedling DNase-seq dataset, of which 65,634 overlapped with those detected by the DNase-seq assay (Figure 4A). Although the remaining 87,960 predicted OCRs were not supported by DNase-seq, 21,420 of these were supported by the *O. sativa* root ATAC-seq dataset (Figure 4B). In Figure 4B, the ATAC-seq data included those of GSE101482 and GSE75794. Based on the currently available DNase-seq and ATAC-seq data of a few plant species, it is reasonable to speculate that more predicted OCRs could be confirmed if more experimental data were available. Among the OCRs predicted in the *O. sativa* seedling data, a considerable proportion was supported by the root data, implying that the OCRs predicted by CharPlant are credible and not false positives.

**Figure 4.**
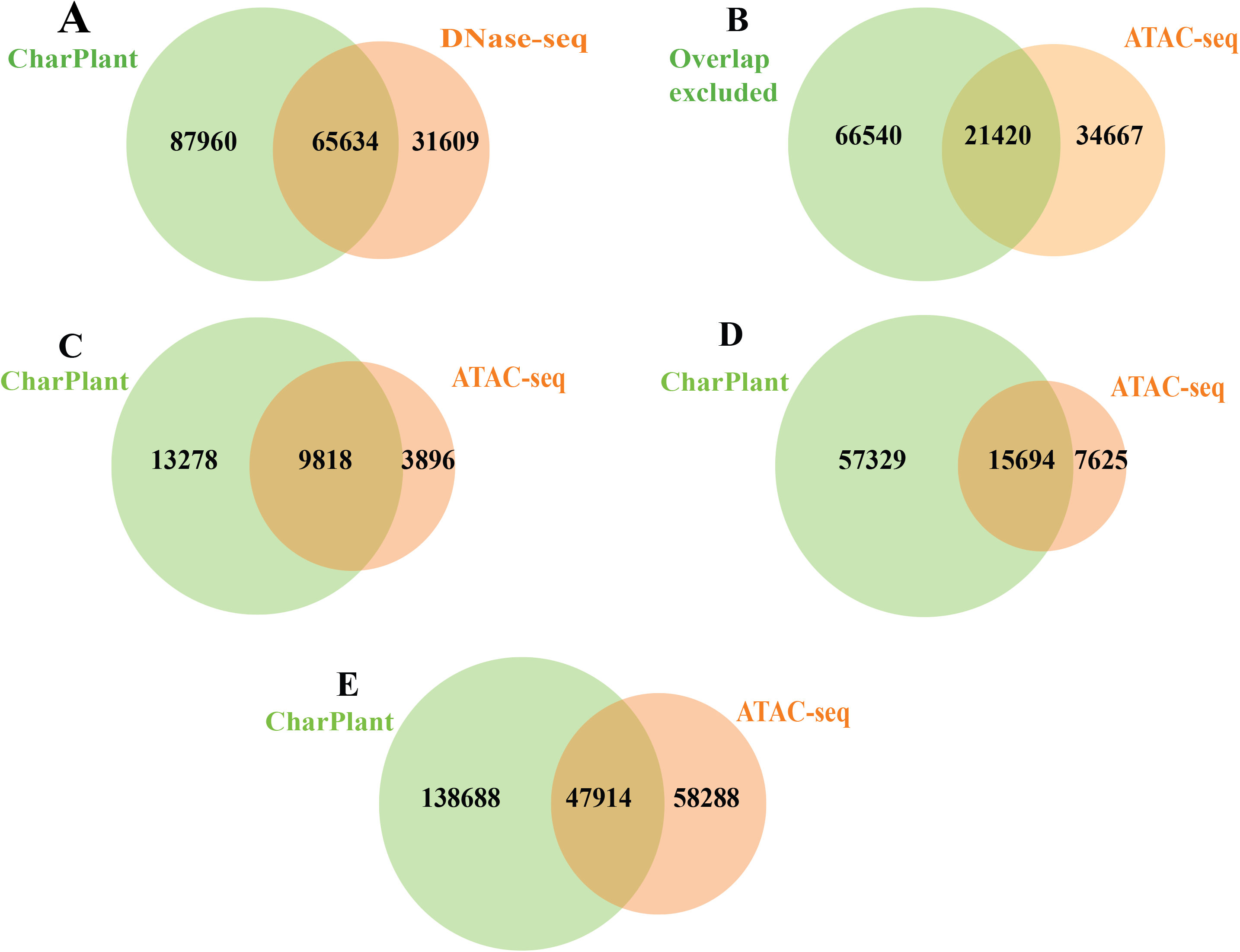
Overlap between the OCRs predicted by CharPlant and those detected by DNase-seq or ATAC-seq assays in four plant species. **A.** Overlap between the predicted OCRs and those detected by DNase-seq in *Oryza sativa*. **B.** Predicted OCRs supported by ATAC-seq after excluding the overlap with DNase-seq. **C.** *Arabidopsis thaliana*. **D.** *Medicago truncatula*. **E.** *Solanum lycopersicum*.

To provide more evidence, we compared the predicted OCRs with experimental OCRs and three histone modifications (H3K4me3, H3K9ac, and H3K27ac) in *A. thaliana*. Covalent modification of the histone tail plays a key role in regulating chromatin structure and gene transcription. In eukaryotes, H3K4me3 is associated with active chromatin and promotes transcription through interactions with effector proteins [41,42]. H3K9ac and H3K4me3 frequently coexist as markers of active gene promoters. H3K27ac is related to gene activation and is mainly enriched in enhancer and promoter regions [43,44]. Figure S4A and B show two examples of overlap between predicted OCRs and experimental OCRs, indicating that the ATAC-seq data supported the predicted results. Additionally, H3K4me3, H3K9ac, and H3K27ac modifications showed peaks at the sites. However, another scenario is where a predicted OCR does not overlap with DNase-seq or ATAC-seq data. For example, in Figure S4C, the ATAC-seq data showed no peak at the predicted OCR, whereas H3K4me3, H3K9ac, and H3K27ac modifications showed significant peaks. Considering the tissue- and time-specificity of OCRs, it was difficult to definitively conclude that this site was not an OCR. To determine whether there was a prediction bias, i.e., some regions have higher prediction accuracy than other regions, we calculated the distribution of predicted OCRs in the promoter, intergenic region, exon, intron, 5’UTR, 3’UTR, and downstream region, and compared them with experimental data to show their consistency. The results showed that the distributions of predicted OCRs were consistent with those of DNase-seq and ATAC-seq data of *O. sativa*, *A. thaliana*, and *M. truncatula* (Figure 5**A-C**). In Figure 5A, the left pie chart was obtained using the DNase-seq data of *O. sativa*. Although the pie charts of *S. lycopersicum* appeared different, the number of OCRs was the highest in the intergenic region, followed by the promoter, and least in the 3’UTR (Figure 5D). Epigenetic modifications provide further evidence for the validation of OCRs predicted by CharPlant. Among the four plant species, *A. thaliana* has the most abundant data, including ATAC-seq dataset and various epigenetic datasets. Therefore, we used *A. thaliana* as an example to calculate the difference in the frequency of the H3K4me3 modification between the predicted OCRs and ATAC-seq peaks (Figure S5A). The distribution of H3K4me3 in the *A. thaliana* genome was obtained from the PCSD database at http://systemsbiology.cau.edu.cn/chromstates [45]. Our results showed no significant difference in the frequency of the H3K4me3 modification between the predicted OCRs and ATAC-seq peaks, and the two boxplots were almost identical. Furthermore, we investigated the difference of H3K4me3 modification between the predicted OCRs and 10,000 randomly selected inactive chromatin regions, based on ATAC-seq data. We found that the predicted OCRs were significantly more enriched in the H3K4me3 modification than the unopened chromatin regions (Figure S5B).

**Figure 5.**
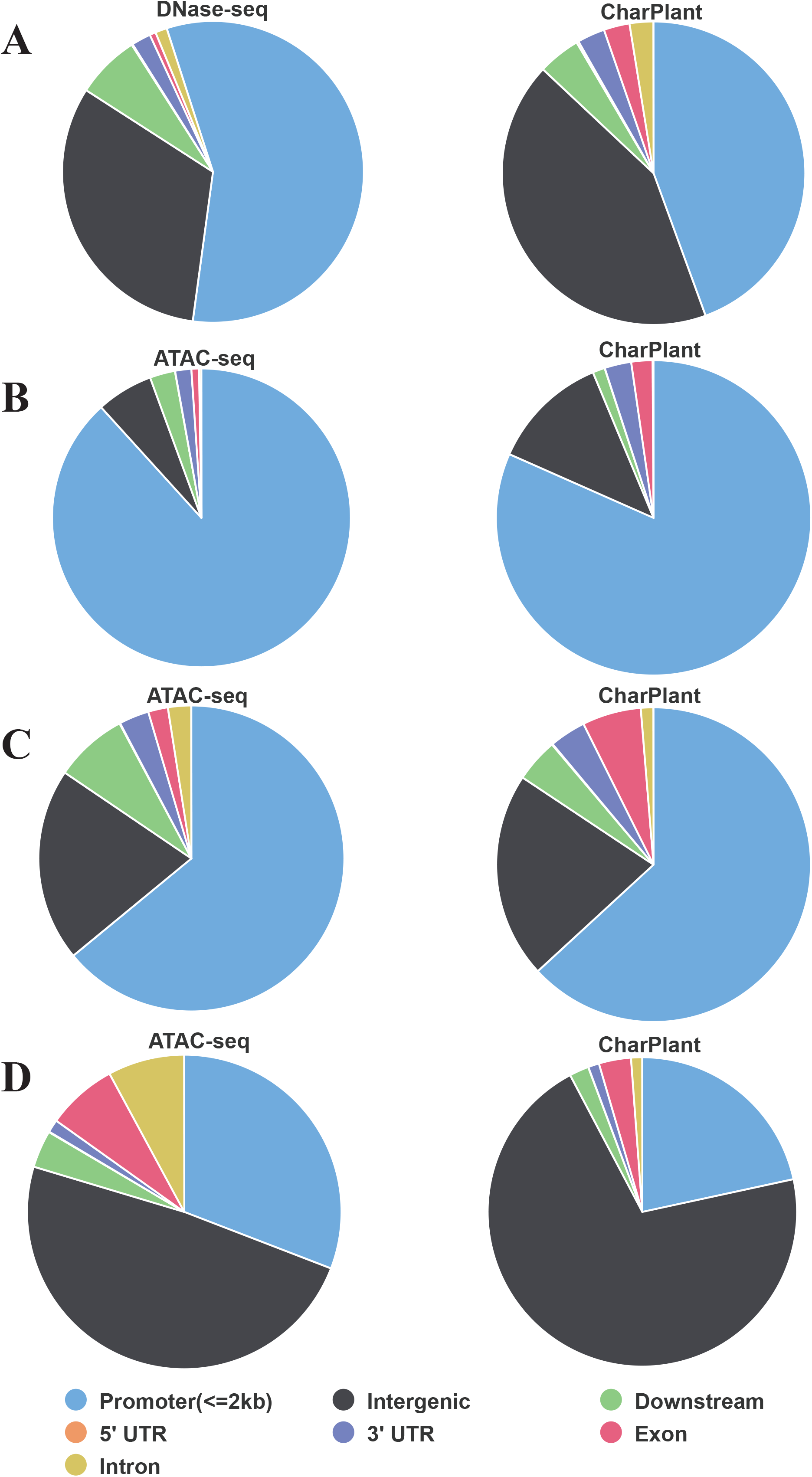
Distribution of predicted OCRs and experimental OCRs in the promoter region, intergenic region, exon, intron, 5’UTR, 3’UTR, and downstream region. **A.** *Oryza sativa*. **B.** *Arabidopsis thaliana*. **C.** *Medicago truncatula*. **D.** *Solanum lycopersicum*.

Notably, the time taken to scan the genome of four plant species was closely related to the genome size; analysis of *A. thaliana*, *O. sativa*, *M. truncatula*, and *S. lycopersicum* genomes took 8, 22, 24, and 49 h, respectively.

## 4 Conclusion

In summary, experimental technologies can determine only the current status of DNA accessibility, whereas CharPlant is a neural network model that learns the sequence motifs and regulatory logic, and predicts potential OCRs, according to the experimental data. Compared with existing algorithms, CharPlant has several advantages. First, to our knowledge, CharPlant is the first *de novo* prediction tool that can identify potential chromatin accessible regions along the genome sequence. Second, CharPlant is specifically designed to predict OCRs of plants, rather than those of human or animals, as in other algorithms. Third, CharPlant marks all potential OCRs of a given plant species in different tissues and at different times, which is beneficial for the investigation of gene regulation under different conditions. Lastly, CharPlant is significantly faster than other deep learning algorithms because it is designed with a concise and efficient structure.

## Data availability

CharPlant is freely available from GitHub at https://github.com/Yin-Shen/CharPlant.

## Authors’ contributions

JG and LLC conceived and designed the project. YS performed computational analysis. JG wrote the first draft of the manuscript. All authors read and approved the final manuscript.

## Competing interests

The authors have declared no competing interests.

## Acknowledgments

This work was supported by the National Natural Science Foundation of China (31871269, 31571351), Hubei Provincial Natural Science Foundation of China (2019CFA014), and Fundamental Research Funds for the Central Universities (2662019PY069).

## Supplementary material

Supplementary material associated with this article can be found in the online version.

## Supplementary material

**Figure S1 ROCs of *Oryza sativa* ATAC-seq data**

ROC, receiver operating curve.

**Figure S2 Precision recall curves of comparative methods in four plant species**

**A**. *Oryza sativa*. **B**. *Arabidopsis thaliana*. **C**. *Medicago truncatula*. **D**. *Solanum lycopersicum.* PR, precision recall.

**Figure S3 Receiver operating curves and precision recall curves of 10-fold cross validation**

**A.** Receiver operating curves. **B**. Precision recall curves. ROCs, receiver operating curves; PR, precision recall.

**Figure S4 Comparison of predicted OCRs, experimental OCRs, and histone modifications H3K4me3, H3K9ac, and H3K27ac in Arabidopsis thaliana**

**A, B**. Two examples showing overlap between predicted OCRs and experimental OCRs.

**C**. An example where ATAC-seq data showed no peak at the predicted OCR, but H3K4me3, H3K9ac, and H3K27ac modifications showed significant peaks.

**Figure S5 Difference in the epigenetic modification H3K4me3 in *Arabidopsis thaliana***

**A.** Between the open chromatin regions predicted by CharPlant and ATAC-seq peaks.

**B.** Between the open chromatin regions predicted by CharPlant and randomly selected inactive regions.

**Supplementary File 1 Steps involved in the installation and execution of CharPlant, config.yaml file, and Snakemake file**.

## References

[1] Klemm SL, Shipony Z, Greenleaf WJ. Chromatin accessibility and the regulatory epigenome. Nat Rev Genet 2019;20:207–20.

[2] Shashikant T, Ettensohn CA. Genome-wide analysis of chromatin accessibility using ATAC-seq. Methods Cell Biol 2019;151:219–35.

[3] Tsompana M, Buck MJ. Chromatin accessibility: a window into the genome. Epigenetics Chromatin 2014;7:33–48.

[4] Thurman RE, Rynes E, Humbert R, Vierstra J, Maurano MT, Haugen E, et al. The accessible chromatin landscape of the human genome. Nature 2012;489:75–82.

[5] Buenrostro JD, Wu B, Litzenburger UM, Ruff D, Gonzales ML, Snyder MP, Chang HY, Greenleaf WJ. Single-cell chromatin accessibility reveals principles of regulatory variation. Nature 2015;523:486–90.

[6] Denny SK, Yang D, Chuang CH, Brady JJ, Lim JS, Gruner BM et al. NFIB promotes metastasis through a widespread increase in chromatin accessibility. Cell 2016;166:328–42.

[7] Osmanbeyoglu HU, Shimizu F, Rynne-Vidal A, Alonso-Curbelo D, Chen HA, Wen HY, et al. Chromatin-informed inference of transcriptional programs in gynecologic and basal breast cancers. Nat Commun 2019;10:1–12.

[8] Qu K, Zaba LC, Satpathy AT, Giresi PG, Li R, Jin Y, et al. Chromatin accessibility landscape of cutaneous T cell lymphoma and dynamic response to HDAC inhibitors. Cancer Cell 2017;32:27–41.

[9] Buenrostro JD, Wu B, Chang HY, Greenleaf WJ. ATAC-seq: A method for assaying chromatin accessibility genome-wide. Curr Protoc Mol Biol 2015;109:21–9.

[10] Song L, Crawford GE. DNase-seq: a high-resolution technique for mapping active gene regulatory elements across the genome from mammalian cells. Cold Spring Harb Protoc 2010.

[11] He HH, Meyer CA, Hu SS, Chen MW, Zang C, Liu Y, et al. Refined DNase-seq protocol and data analysis reveals intrinsic bias in transcription factor footprint identification. Nat Methods 2014;11:73–78.

[12] Sung MH, Guertin MJ, Baek S, Hager GL. DNase footprint signatures are dictated by factor dynamics and DNA sequence. Mol Cell 2014;56:275–85.

[13] Lu Z, Hofmeister BT, Vollmers C, Dubois RM, Schmitz RJ. Combining ATAC-seq with nuclei sorting for discovery of cis-regulatory regions in plant genomes. Nucleic Acids Res 2017;45:e41.

[14] Milan M, Balestrieri C, Alfarano G, Polletti S, Prosperini E, Spaggiari P, et al. FOXA2 controls the cis-regulatory networks of pancreatic cancer cells in a differentiation grade-specific manner. Embo J 2019:e102161.

[15] Zhang T, Marand AP, Jiang J. PlantDHS: a database for DNase I hypersensitive sites in plants. Nucleic Acids Res 2016;44:D1148–53.

[16] Maher KA, Bajic M, Kajala K, Reynoso M, Pauluzzi G, West DA, et al. Profiling of accessible chromatin regions across multiple plant species and cell types reveals common gene regulatory principles and new control modules. Plant Cell 2018;30:15–36.

[17] Steinmuller K, Batschauer A, Apel K. Tissue-specific and light-dependent changes of chromatin organization in barley (Hordeum vulgare). Eur J Biochem 1986;158:519–25.

[18] Fang S, Li J, Xiao Y, Lee M, Guo L, Han W, et al. Tet inactivation disrupts YY1 binding and long-range chromatin interactions during embryonic heart development. Nat Commun 2019;10:4297.

[19] Gao L, Wu K, Liu Z, Yao X, Yuan S, Tao W, et al. Chromatin accessibility landscape in human early embryos and its association with evolution. Cell 2018;173:248–59.

[20] Sen DR, Kaminski J, Barnitz RA, Kurachi M, Gerdemann U, Yates KB, et al. The epigenetic landscape of T cell exhaustion. Science 2016;354:1165–9.

[21] Cusanovich DA, Reddington JP, Garfield DA, Daza RM, Aghamirzaie D, Marco-Ferreres R, et al. The cis-regulatory dynamics of embryonic development at single-cell resolution. Nature 2018;555:538–42.

[22] Zhang W, Wu Y, Schnable JC, Zeng Z, Freeling M, Crawford GE, et al. High-resolution mapping of open chromatin in the rice genome. Genome Res 2012;22:151–62.

[23] Zhang W, Zhang T, Wu Y, Jiang J. Genome-wide identification of regulatory DNA elements and protein-binding footprints using signatures of open chromatin in *Arabidopsis*. Plant Cell 2012;24:2719–31.

[24] Voss TC, Hager GL. Dynamic regulation of transcriptional states by chromatin and transcription factors. Nat Rev Genet 2014;15:69–81.

[25] Kelley DR, Snoek J, Rinn JL. Basset: learning the regulatory code of the accessible genome with deep convolutional neural networks. Genome Res 2016;26:990–9.

[26] Chen KM, Cofer EM, Zhou J, Troyanskaya OG. Selene: a PyTorch-based deep learning library for sequence data. Nat Methods 2019;16:315–8.

[27] Alipanahi B, Delong A, Weirauch MT, Frey BJ. Predicting the sequence specificities of DNA- and RNA-binding proteins by deep learning. Nat Biotechnol 2015;33:831–8.

[28] Angermueller C, Lee HJ, Reik W, Stegle O. DeepCpG: accurate prediction of single-cell DNA methylation states using deep learning. Genome Biol 2017;18:67–79.

[29] Mainiero S, Pawlowski WP. Meiotic chromosome structure and function in plants. Cytogenetic & Genome Research 2014;143:6–17.

[30] Zhang Y, Liu T, Meyer CA, Eeckhoute J, Johnson DS, Bernstein BE, et al. Model-based analysis of ChIP-Seq (MACS). Genome Biol 2008;9:R137.

[31] Bailey TL, Boden M, Buske FA, Frith M, Grant CE, Clementi L, et al. MEME SUITE: tools for motif discovery and searching. Nucleic Acids Res 2009;37:W202–8.

[32] Krizhevsky A, Sutskever I, Hinton GE. ImageNet classification with deep convolutional neural networks. Advances in Neural Information Processing Systems 2012;25:1–9.

[33] Lecun Y, Bengio Y, Hinton G. Deep learning. Nature 2015;521:436–44.

[34] Ge W. A perspective on deep imaging. IEEE Access 2017, 4:8914–24.

[35] Young T, Hazarika D, Poria S, Cambria E. Recent trends in deep learning based natural language processing. IEEE Computational Intelligence Magazine 2018;13:55–75.

[36] Bengio Y, Courville A, Vincent P. Representation Learning: A review and new perspectives. IEEE Transactions on Pattern Analysis and Machine Intelligence 2013;35:1798–828.

[37] Mathelier A, Zhao X, Zhang AW, Parcy F, Worsley-Hunt R, Arenillas DJ, et al. JASPAR 2014: an extensively expanded and updated open-access database of transcription factor binding profiles. Nucleic Acids Res 2014;42:D142–7.

[38] Köster J, Rahmann S. Snakemake-a scalable bioinformatics workflow engine. Bioinformatics 2012;28:2520–2.

[39] Hashim FA, Mabrouk MS, Al-Atabany W. Review of different sequence motif finding algorithms. Avicenna J Med Biotechnol 2019;11:130–48.

[40] Pedregosa F, Varoquaux G, Gramfort A, Michel V, Thirion B, Grisel O, et al. Scikit-learn: machine learning in python. Journal of Machine Learning Research 2011;12:2825–30.

[41] Pena PV, Davrazou F, Shi X, Walter KL, Verkhusha VV, Gozani O, et al. Molecular mechanism of histone H3K4me3 recognition by plant homeodomain of ING2. Nature 2006;442:100–3.

[42] Schneider R, Bannister AJ, Myers FA, Thorne AW, Crane-Robinson C, Kouzarides T. Histone H3 lysine 4 methylation patterns in higher eukaryotic genes. Nat Cell Biol 2004;6:73–7.

[43] Musselman CA, Lalonde ME, Cote J, Kutateladze TG. Perceiving the epigenetic landscape through histone readers. Nat Struct Mol Biol 2012;19:1218–27.

[44] Sproul D, Gilbert N, Bickmore WA. The role of chromatin structure in regulating the expression of clustered genes. Nat Rev Genet 2005;6:775–81.

[45] Liu Y, Tian T, Zhang K, You Q, Yan H, Zhao N, et al. PCSD: a plant chromatin state database. Nucleic Acids Res 2018;46:D1157–67.

