## Supplementary figures and images for "CharPlant: A *De Novo* Open Chromatin Region (OCR) Prediction Tool for Plant Genomes"

### Figure S1

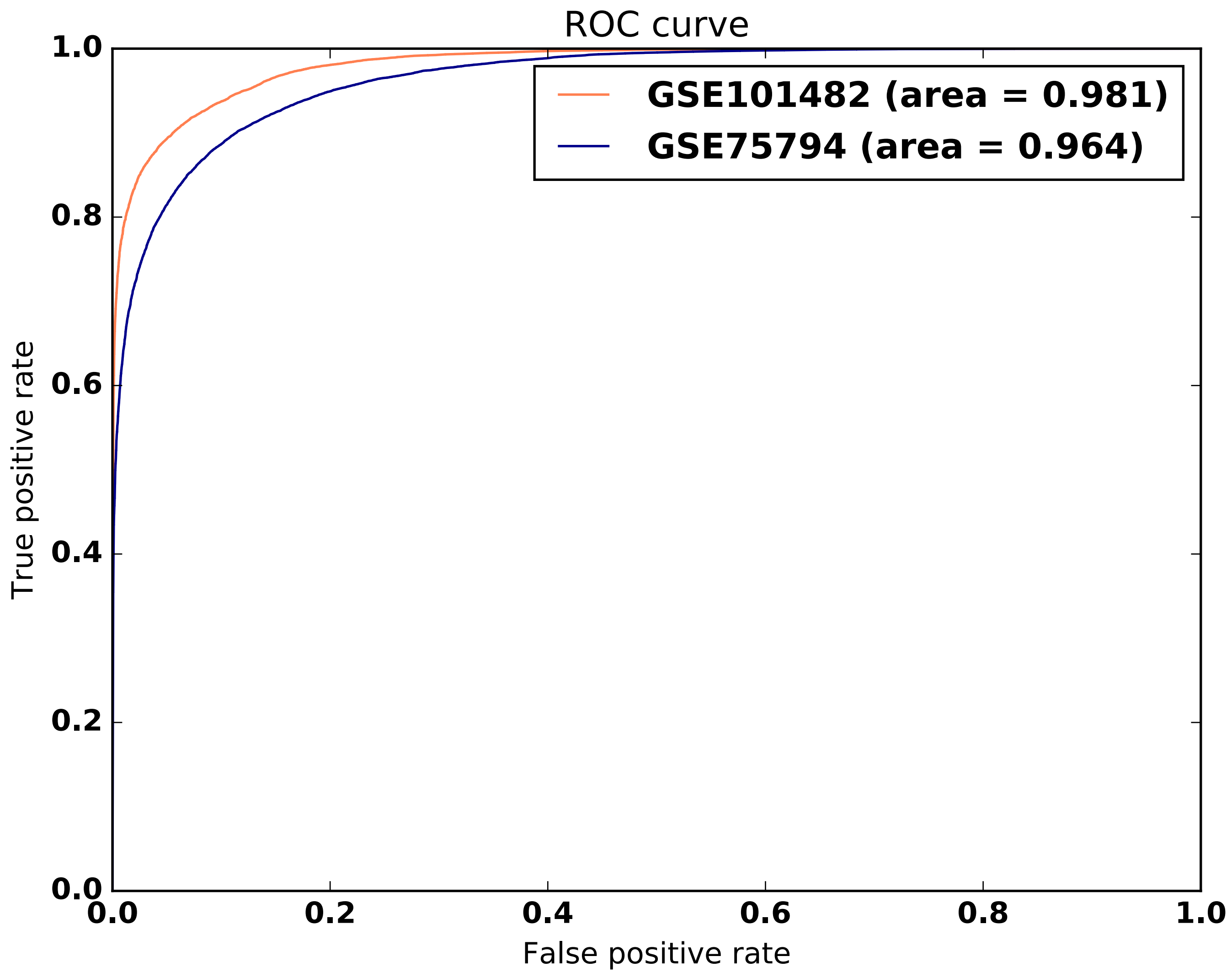

### Figure S2

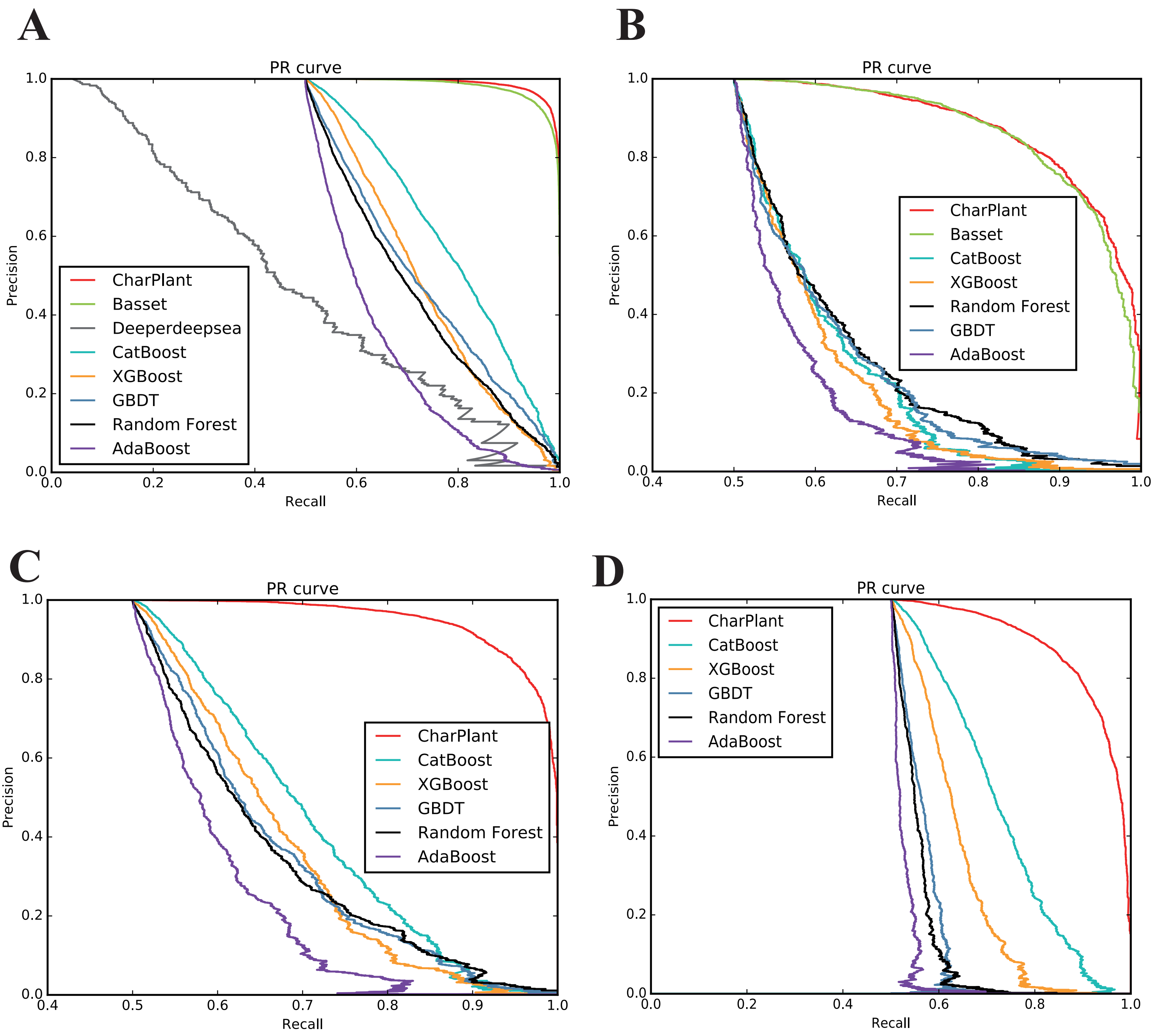

### Figure S3

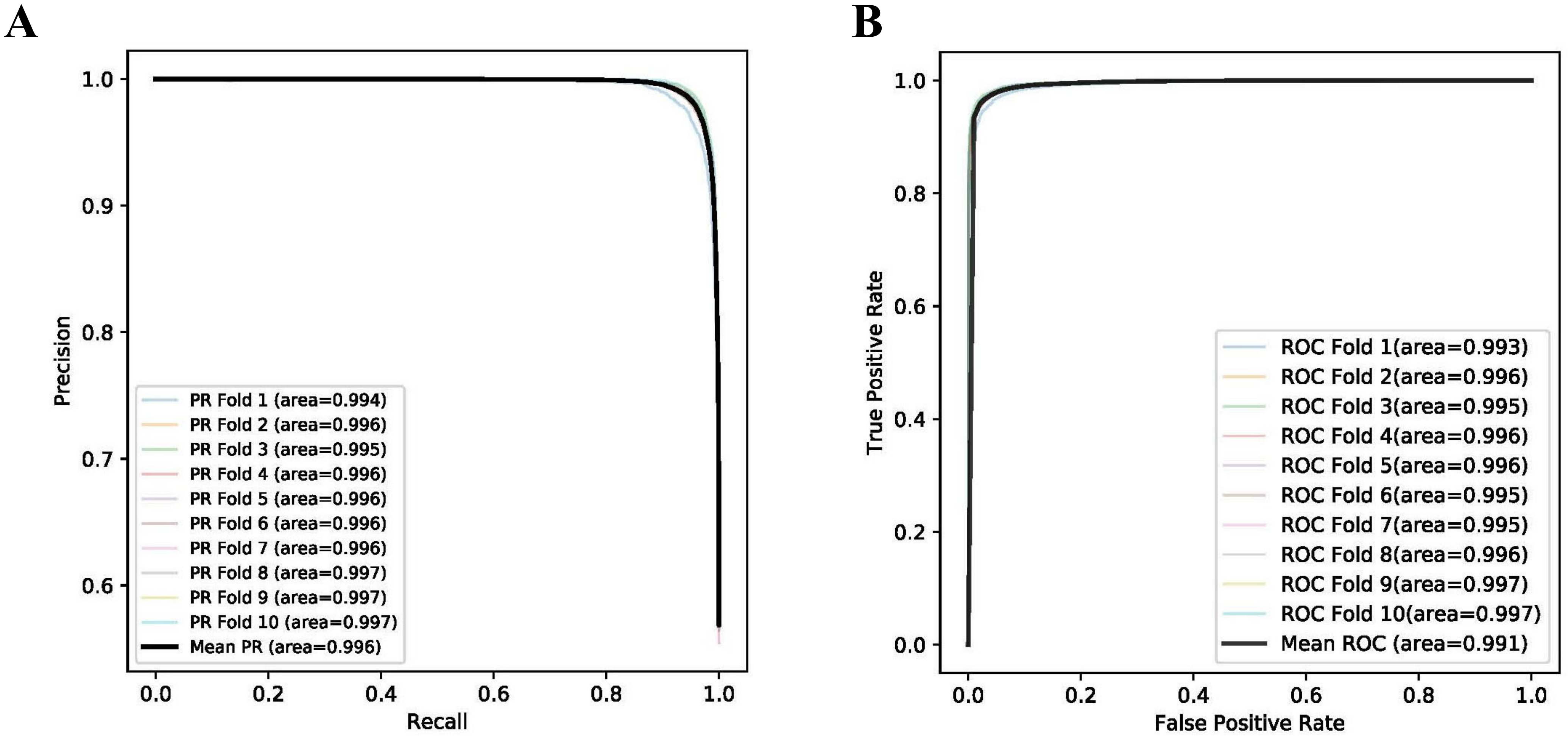

### Figure S4

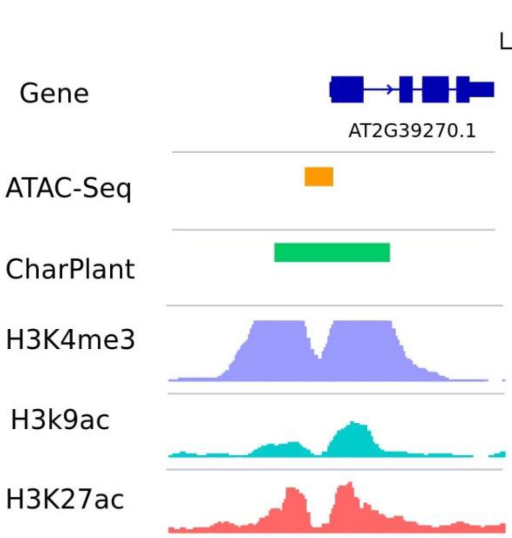

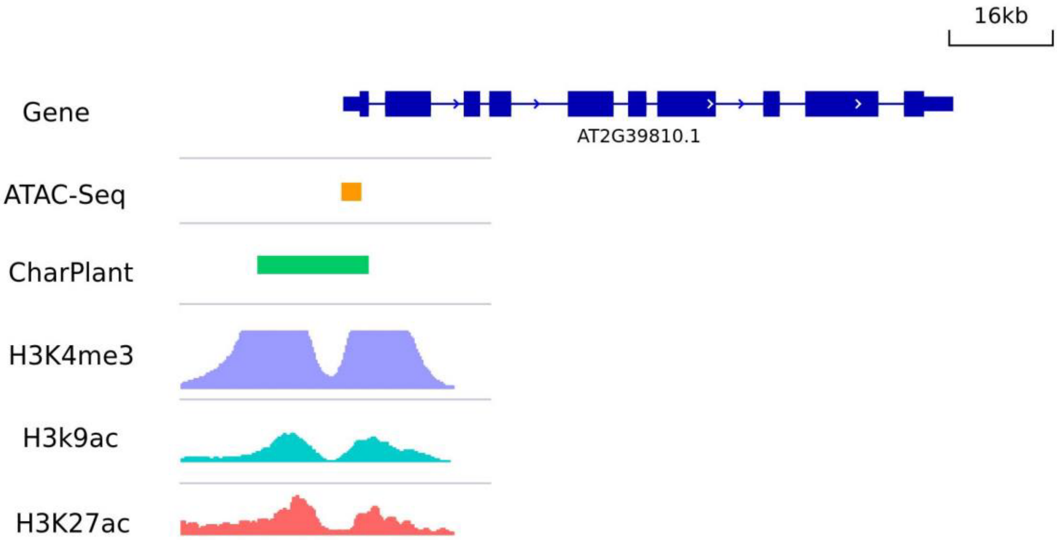

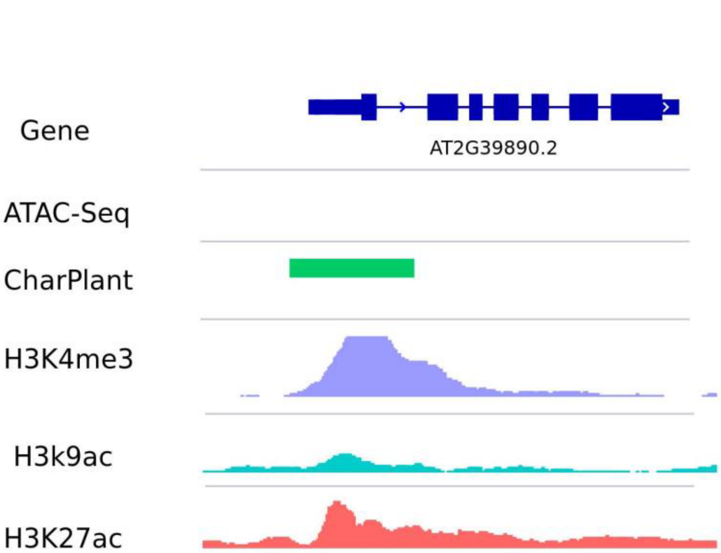

### Figure S5

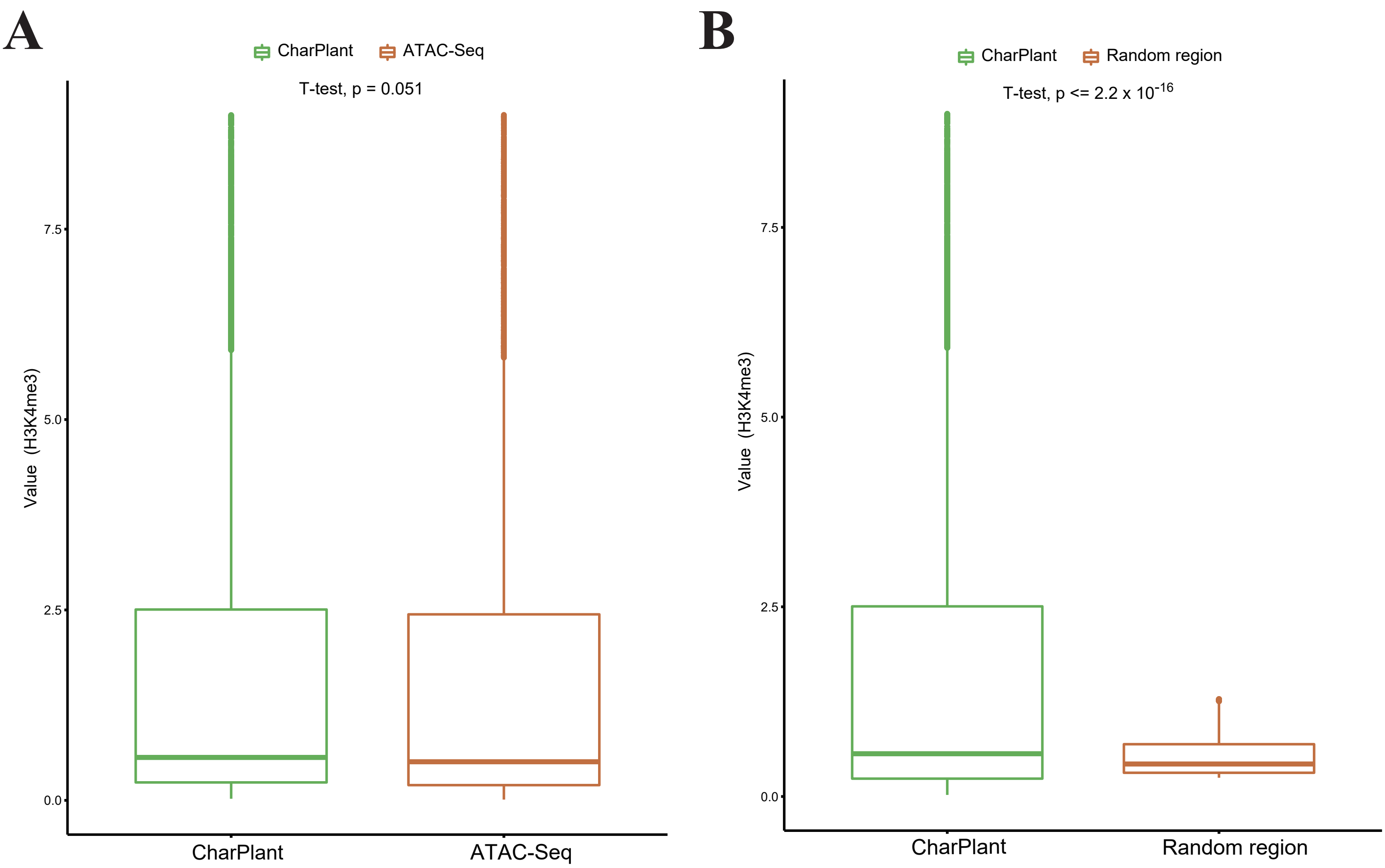
